# A bacterial size law revealed by a coarse-grained model of cell physiology

**DOI:** 10.1101/078998

**Authors:** François Bertaux, Julius von Kügelgen, Samuel Marguerat, Vahid Shahrezaei

## Abstract

Universal observations in Biology are sometimes described as “laws”. In *E. coli*, experimental studies performed over the past six decades have revealed major growth laws relating ribosomal mass fraction and cell size to the growth rate. Because they formalize complex emerging principles in biology, growth laws have been instrumental in shaping our understanding of bacterial physiology. Here, we discovered a novel size law that connects cell size to the inverse of the metabolic proteome mass fraction and the active fraction of ribosomes. We used a simple whole-cell coarse-grained model of cell physiology that combines the proteome allocation theory and the structural model of cell division. The model captures all available experimental data connecting the cell proteome composition, ribosome activity, division size and growth rate in response to nutrient quality, antibiotic treatment and increased protein burden. Finally, a stochastic extension of the model explains non-trivial correlations observed in single cell experiments including the adder principle. This work provides a simple and robust theoretical framework for studying the fundamental principles of cell size determination in unicellular organisms.

## Introduction

The behavior of complex biological systems can be described by surprisingly simple rules or “laws”, which connect quantitatively aspects of the cell composition with its physiology. Over the last six decades, the discovery of several rules, called the bacterial growth laws, transformed the field of microbial physiology (Schaechter et al., 1958; Schleif, 1967; Cooper and Helmstetter, 1968; Maaløe, 1969; Churchward et al., 1982; Klumpp and Hwa, 2014; Vadia and Levin, 2015; Jun et al., 2018). Growth laws describe relationships between the exponential growth rate and cellular parameters. One such law states that the proportion of the cellular content dedicated to ribosomes is linearly dependent on the growth rate (we call this hereafter the first growth law). A second growth law states that average cell size is an exponential function of the growth rate (referred to hereafter as the second growth law). These laws formalize emerging relations between cellular processes quantitatively and therefore generate testable hypothesis for mechanistic studies of the molecular circuitry that underpin cell physiology (Hill et al., 2013; Scott et al., 2014).

What are the biological principles underlying the first growth law? The law was first rationalized as a direct consequence of the catalytic role of ribosomes in protein synthesis (Neidhardt and Magasanik, 1960). Indeed, if every ribosome in a cell synthesises proteins at a constant condition-independent rate, then the steady-state growth rate will be proportional to the fraction of cellular content devoted to ribosomes. However, this does not explain what sets the amount of resources invested into ribosomes in a given environment, and what prevents the cell to increase this investment in order to grow faster. Theoretical and experimental studies identified cellular resource allocation as a key concept to address these questions (Basan, 2018). The theory of resource allocation describes how different groups of proteins with similar function (proteome sectors) are regulated reciprocally in conditions that affect growth rate. In particular, these studies showed that cells invest higher amounts into ribosomes when nutrient quality increases because the investment in metabolic proteins required to achieve a given growth rate is reduced. This explains the positive correlation of the proteome fraction dedicated to ribosomal proteins with the growth rate and provides a model for how the fraction of ribosomes is set. Interestingly, Hwa and co-workers (Scott et al., 2010) also showed that the first growth law is not always valid when growth rate is not modulated by nutrient quality. While the law remains valid when useless proteins are over-expressed, it breaks down when translation is inhibited. Based on these findings, they proposed a phenomenological model of proteome allocation whose predictions go beyond the first growth law because it captures these orthogonal types of growth rate modulation (Scott et al., 2010). More mechanistic coarse-grained models predicting both the growth rate and the coarse-grained proteome as a function of growth conditions have also been proposed confirming and extending these findings (Marr, 1991; Molenaar et al., 2009; Weiße et al., 2015; Maitra and Dill, 2015; Bosdriesz et al., 2015; Maitra and Dill, 2016; Pandey and Jain, 2016; Giordano et al., 2016; Liao et al., 2017; Thomas et al., 2018; Sharma et al., 2018; Pandey et al., 2018).

In contrast, the mechanisms underlying the second growth law, which connects growth rate and cell size, are less well understood. In 1968, Donachie (Donachie, 1968) proposed that, in bacteria, an exponential dependency of cell size on growth rate indicates that: 1) the timing of DNA replication is fully controlled by cell size, and is triggered at a constant volume per origin of replication; 2) cell division is enslaved to the DNA replication processes, and occurs after a constant time-interval following replication initiation, which encompasses DNA replication and cell division (*C+D* period). However, while recent work has confirmed that DNA replication is strongly size-controlled (Wallden et al., 2016) and that the initiation size per origin is invariant across a large range of growth rates and types of growth rate modulations (Si et al., 2017), what determines the duration of the *C+D* period is still unclear. Moreover, the relationship between cell size and growth rate does not follow the second growth law, when useless proteins are over-expressed on gene expression is increased or translation is inhibited (Basan et al., 2015; Si et al., 2017). This indicates that duration of the *C+D* period is regulated in non-trivial ways outside of the canonical growth modulations based on nutrient quality. To date, unlike for the first growth law, mechanistic understanding of this phenomenon in connection to the second growth law is lacking.

The two growth laws discussed so far are based on average cell population measurements. However, the variations at the single cell level from the population average also provides rich and complementary insights into the biology of cell growth and division (Taheri-Araghi et al., 2015; Kennard et al., 2016). Cells can control their size through a ‘sizer’ mechanism, where regardless of birth size they grow to an average size and then they divide. But, recent advances in single-cell techniques led to the discovery that bacterial cells achieve cell size homeostasis via an ‘adder’ principle, where cells add a constant size between birth and division independent of the birth size (Campos et al., 2014; Taheri-Araghi et al., 2015; Amir, 2014). The adder phenomenon has been observed in several bacteria, yeast, archea and mammalian cells (Iyer-Biswas et al., 2014; Taheri-Araghi et al., 2015; Deforet et al., 2015; Soifer et al., 2016; Chandler-Brown et al., 2017; Logsdon et al., 2017; Priestman et al., 2017; Varsano et al., 2017; Eun et al., 2018; Cadart et al., 2018). Despite its broad conservation, the mechanistic basis of this phenomenon remains elusive and hotly debated. It has been suggested that the adder phenomenon is related to second growth law assuming that replication controls cell division (Ho and Amir, 2015; Amir, 2017). However, this model is challenged by recent studies that questioned the role of DNA replication in controlling division of single cells (Harris and Theriot, 2016; Micali et al., 2018; Si et al., 2019).

Here, we present a coarse-grained model of bacterial physiology that unifies the proteome allocation theory with the structural model of division control (Basan et al., 2015; Fantes et al., 1975; Taheri-Araghi et al., 2015). With a small number of fitting parameters, the model captures a wide range of experimental measurements of protein mass fraction, ribosome activity and cell size across growth conditions revealing a novel bacterial size law where cell size depends on the metabolism proteome sector mass fraction and the fraction of active ribosomes. This new law is simple and more general than the original second growth law. Finally, the model exhibits single cell properties that are consistent with experimental observations including the adder size control.

## Results

### A minimalistic whole-cell coarse-grained model to predict cell composition, growth rate and cell size

We developed a minimalistic whole-cell coarse-grained mathematical model to predict cell composition, growth rate and cell size for at least three types of growth rate modulation: 1) change of nutrient quality, 2) translation inhibition by chloramphenicol and 3) over-expression of useless proteins. These modulations were selected because they have been instrumental in uncovering key principles of proteome allocation underpinning the first growth law (Scott et al., 2010) and have abundant experimental measurements available (Scott et al., 2010; Basan et al., 2015; Taheri-Araghi et al., 2015; Dai et al., 2017; Si et al., 2017). Our model extends the existing whole-cell coarse-grained models (Marr, 1991; Molenaar et al., 2009; Weiße et al., 2015; Maitra and Dill, 2015; Bosdriesz et al., 2015; Maitra and Dill, 2016; Pandey and Jain, 2016; Giordano et al., 2016; Liao et al., 2017; Thomas et al., 2018; Sharma et al., 2018; Pandey et al., 2018) by including both cell composition and cell size and capturing data from several studies: size data (Basan et al., 2015; Taheri-Araghi et al., 2015; Si et al., 2017), proteome allocation data (Scott et al., 2010; Dai et al., 2017) and ribosome activity data (Dai et al., 2017).

As in previous work, the model (Figure 1, see Methods for a mathematical description) considers only two types of molecular components: protein precursors (noted *A*) and proteins (Molenaar et al. 2009; Weisse et al. 2015; Pandey & Jain 2016). Proteins are split between four different classes (or sectors): transport and metabolism proteins (referred to as only metabolic sector for brevity) (*E*), ribosomal proteins (R), housekeeping proteins (*Q*), and division proteins (*X*) (Basan et al., 2015; Taheri-Araghi et al., 2015). *E* proteins catalyse the import and transformation of external nutrients (not explicitly modelled) into precursors A. R proteins are involved in the synthesis of proteins from precursors. *Q* proteins are involved in cellular functions that do not directly contribute to growth. *X* proteins are regulating cell division (see below). Lastly, when growth rate is modulated by over-expression of useless proteins, we consider an additional class of “useless” proteins noted *U* (Figure 1). All proteins are stable in our model and only diluted by cell growth. The model enforces mass conservation-cell mass being defined as the sum of all molecular components. Finally, except for the import and transformation of nutrients into precursors, all reactions leave cell mass unchanged and assumes mass density homeostasis (cell volume and cell mass are therefore equivalent), as observed experimentally (Basan et al., 2015).

**Figure 1.**
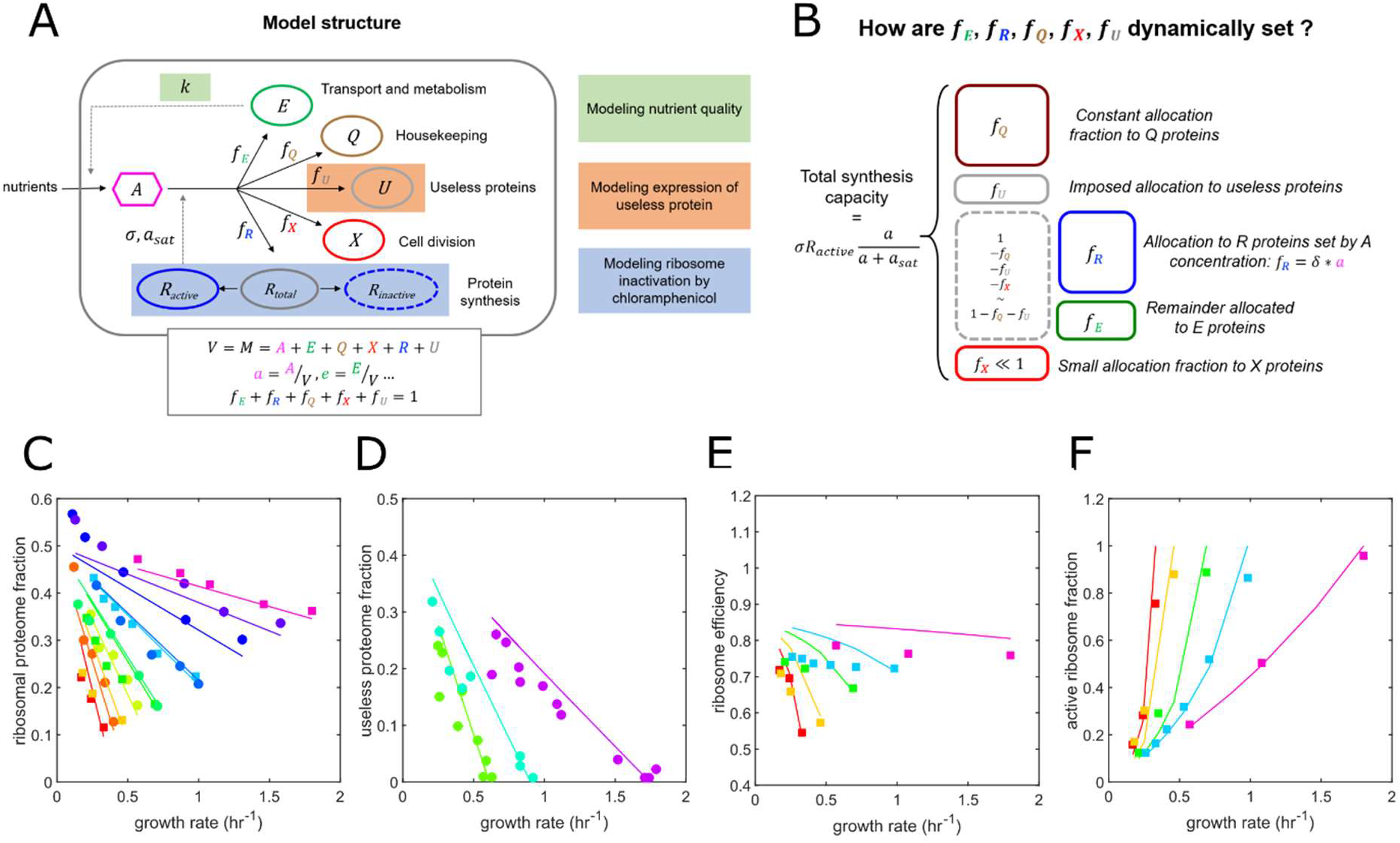
A simple whole-cell coarse-grained model of bacterial growth reproduces proteome allocation and ribosome activity data. (A) Schematic of model structure displaying model components (proteome sectors E, R, Q and X and protein precursor A) and reactions (metabolism, protein synthesis). How the three types of growth rate modulation (nutrient quality, expression of useless protein and ribosome inactivation) are modeled is also highlighted. The rate of precursor creation is proportional to the amount of metabolic enzymes E. The total protein synthesis rate is proportional to the number of active ribosomes, and the synthesis rate per ribosome (the ribosome efficiency) is dependent on the precursor concentration via a Michaelis-Menten relationship. A full description of the model is given in the Methods section. (B) Allocation of total protein synthesis capacity between proteome sectors. A fixed, condition-independent fraction f_Q_ is allocated to housekeeping proteins. Expression of useless protein imposes a fixed allocation f_ν_. The allocation to R proteins is proportional to precursor concentration, and the remainder is allocated to E proteins (assuming that f_X_ ≪ 1). (C-F) Model predictions (solid lines) agree with experimental data (circles – Scott et al. 2010, squares – Dai et al. 2017). Parameters were all fixed from published data (see Methods). (C) Ribosomal proteome fraction data for nutrient and chloramphenicol growth rate modulations. Colors indicate different nutrient qualities. (D) Relationship between growth rate and useless proteome fraction for forced expression of useless protein. (E-F) Ribosome efficiency 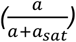 and active ribosome fraction 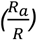.

To fully specify the model, one should describe how the protein synthesis allocation fractions (how much of the total protein synthesis capacity is devoted to each protein class) are set (Figure 1B). A fixed, condition-independent fraction *f_Q_* is allocated to Q proteins. We assume that the allocation fraction to division proteins is small (*f_X_* ≪ 1). The allocation fraction *f_U_* is imposed and represents the level of useless protein expression. The main allocation decision is therefore between *f_R_* and *f_E_*. In this study, rather than assuming growth rate maximization to set these fractions, we assume that *f_R_* (and hence *f_E_*) is dynamically regulated as detailed below. This choice is motivated by the recent observation (Dai et al., 2017) that the ribosome translation elongation rate exhibits a Michaelis-Menten dependence on the total ribosome proteome fraction across nutrient and chloramphenicol-mediated growth modulations. Interestingly, this can be simply reproduced in our model by assuming that the allocation fraction to ribosomal proteins *f_R_* is proportional to the precursor concentration *a*, *f_R_* = *δa* (Methods). We further assume that *δ* is large enough to ensure that *a* ≪ 1, as the steady-state concentration of free amino-acids is known to remain small (Scott et al., 2014). Simulations show that this approximation is already valid when *δ* = 5 (Supplementary Figure 1). Then, the steady-state cell composition and growth rate can be predicted and depend only on: 1) three condition-independent parameters: the maximal ribosome speed *σ,* the ribosome regulation constant *K* and the housekeeping protein allocation fraction *f_Q_*; and 2) three growth modulation parameters: the medium nutrient quality *k,* the chloramphenicol-imposed inactivation rate of ribosomes 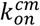, and the useless protein allocation fraction *f_U_*.

The model reproduces quantitatively proteome allocation data (Scott et al., 2010; Dai et al., 2017) and ribosome activity data (Dai et al., 2017) for the three types of growth modulation (Figure 1C-F). This agreement is obtained without parameter fitting, because the parameters *σ*, *K* and *f_Q_* are directly constrained from experimental measurements (Scott et al., 2010; Dai et al., 2017). To our knowledge, this is the first coarse-grained model of bacterial growth to reproduce both proteome allocation and ribosome activity data.

While it is often assumed that proteome fractions are optimally allocated to maximize growth rate (Molenaar et al., 2009), in our model proteome fractions are dynamically set by the simple regulation *f_R_* = *δa*. Interestingly, this regulation achieves near-optimal growth rates across the three types of growth rate modulations considered in this study (Supplementary Figure 2). However, we note that the model assuming optimal proteome allocation does not reproduce the ribosome activity data of Figure 1E.

Our model also explicitly includes cell division and assumes that it is triggered by the accumulation of proteins belonging to the *X* sector to an absolute copy number threshold *X_div_* (Figure 2A), as proposed before in the structural model (Fantes et al., 1975; Taheri-Araghi et al., 2015; Basan et al., 2015). This design enables prediction of changes in both cell size and coarse-grained cell composition as a function of cell growth. Note that for simplicity and because it appears unnecessary, we do not assume that *X* proteins are destroyed immediately after division, in contrast to other studies (Fantes et al., 1975; Taheri-Araghi et al., 2015). Altogether, our model assumptions result in simple and intuitive equations characterizing model steady-states (Figure 2B). Importantly, steady-state concentrations of cell components are not dependent on the cell cycle as expected under balanced growth and are valid when cell division is not modeled (as done for the fitting of the proteome fraction data above).

**Figure 2.**
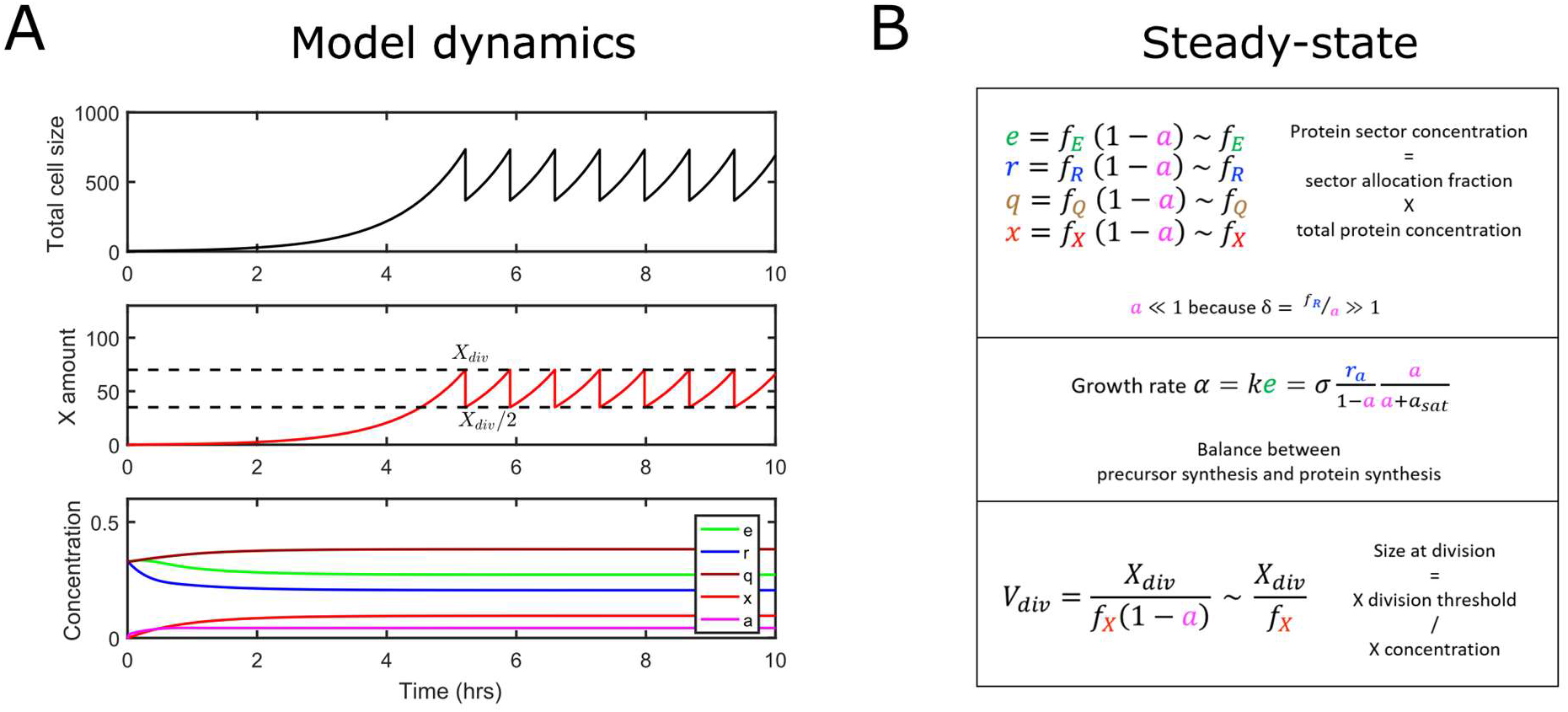
Integration of the structural model enables prediction of both cell composition and cell size. (A) Simulation of model dynamics. After cell division (triggered when X number reaches X_div_), cell content is partitioned equally among daughter cells and only one is followed (‘mother machine’ setting, Wang et al., 2010). After a transient adaptation period, concentrations of cellular components are constant, cellular growth (size increase) is exponential and division occurs at a constant size. Here, nutrient quality is such that the growth rate is 1 hr^−1^ and there is no chloramphenicol nor useless expression. (B) Equations characterizing model steady-state together with their intuitive interpretation (see Methods for their derivation). r_a_ denotes the concentration of active ribosomes.

In summary, we have developed a whole-cell model of cell physiology that captures changes in proteome fractions and ribosome activity as a function of growth conditions while uniquely incorporating direct modeling of cell division control.

### Cell size can be predicted from coarse-grained cell composition across all conditions revealing a novel size law

Experimental data from previous studies shows a complex relationship between cell size and growth rate across the three types of growth modulation (Figure 3B, Supplementary Figure 3). Cell size increases with growth rate when nutrient quality is varied as posited by the second growth law, but the dependency on growth rate is inverted when growth rate is modulated by forced expression of useless proteins. Moreover, chloramphenicol-mediated translation inhibition could increase or decrease cell size depending on the nutrient quality of the medium. We asked whether particular regulation of division proteins *X* by coarse-grained proteome sectors via the allocation fraction *f_X_* could explain this complex pattern (Figure 3A). If the division threshold *X_div_* is invariant, cell size follows the inverse of *f_X_* (Figure 2B, bottom), so regulation of *X* by coarse-grained proteome sectors directly links the cell size to the size of those sectors.

**Figure 3.**
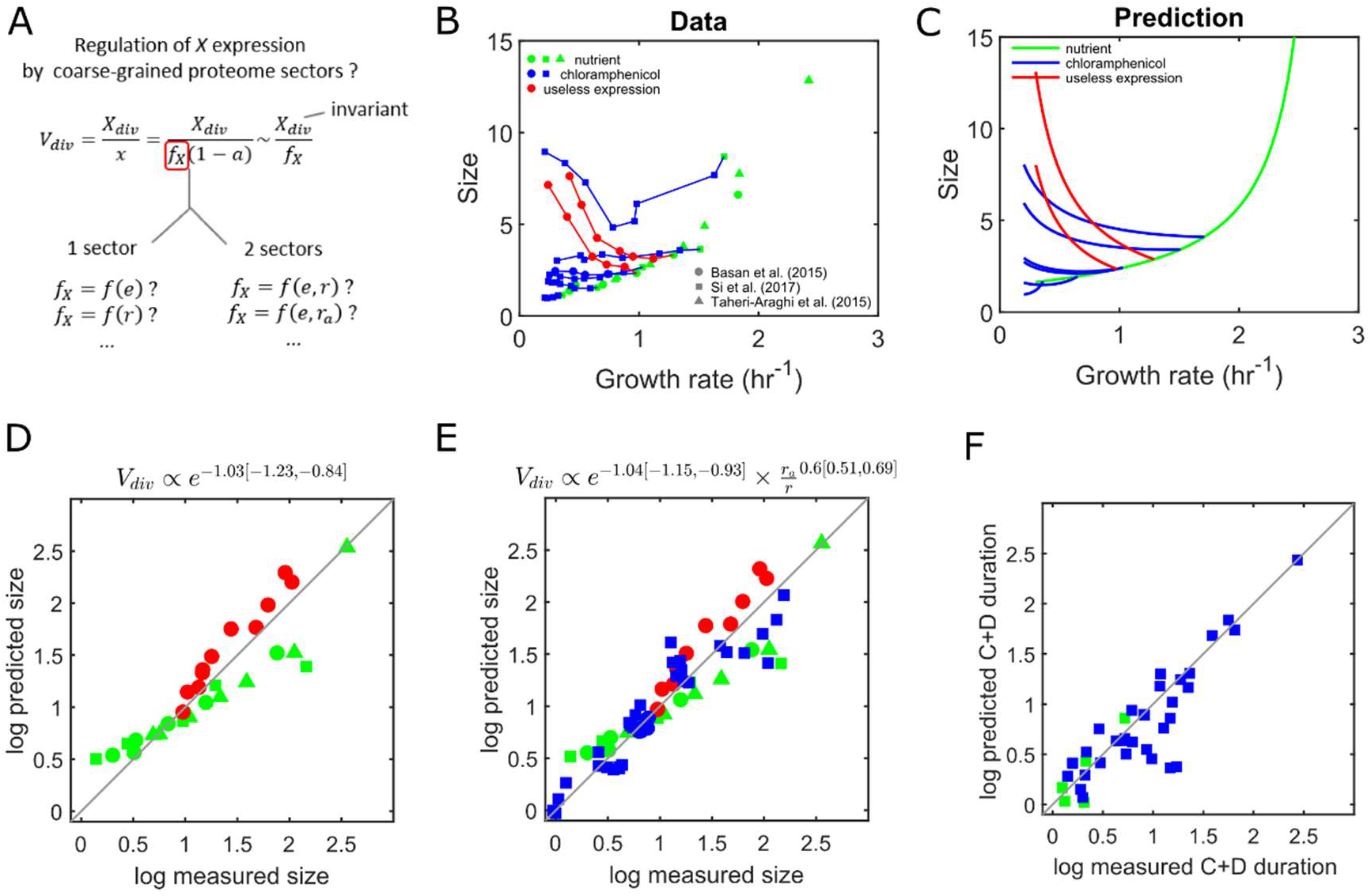
Regulation of division proteins by two proteome sectors quantitatively explain cell size across growth modulations. (A) Hypothesis stating that X expression could depend on the concentration of one or two coarse-grained proteome sectors. Here we assume that X_div_ is invariant. (B) Empirical relationship between cell size and growth rate for three types of growth rate modulation (nutrient quality, chloramphenicol-mediated translation inhibition and expression of useless protein). Data aggregated from three studies, Basan et al. (2015), Si et al. (2017) and Taheri-Araghi et al. (2015). A scaling factor for size was applied on Si et al. and Taheri-Araghi et al. data to make the nutrient modulation data of the three studies consistent (see Supplementary Figure 3). Different branches for useless expression and chloramphenicol indicate modulations at different nutrient quality. (C) Predicted relationship between cell size and growth rate when X expression depends on both E concentration and the fraction of active ribosomes. (D-F) Log-log plots comparing model predictions with experimental data. The natural log is used. (D) Regulation of X expression by E alone can explain size data for nutrient modulation and useless expression modulation (but not chloramphenicol-mediated translation inhibition, Supplementary Figure 4). (E) Among possible regulations of X by two coarse-grained quantities, regulation by E concentration and the fraction of active ribosomes explains size data for all three types of growth rate modulations (this regulation model was used for (C)). (F) Predicted C+D durations agree with measurements by Si et al. (2017). C+D is predicted by 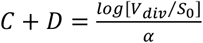, where α is the growth rate, V_div_ is the model-predicted size (3C and 3E) and S_0_ is a constant measured in Si et al. (2017).

We first considered a model whereby *f_X_* follows the concentration of a single coarse-grained proteome sector. Interestingly, size measurements for both nutrient quality and useless protein expression growth rate modulations are explained if *f_X_* is set to follow the concentration (noted *e*, not to confuse with the mathematical constant) of the metabolic sector *E* exclusively (Figure 3D). However, this simple dependency cannot explain chloramphenicol-mediated translation inhibition data (Supplementary Figure 4). We therefore extended this analysis to dependencies of *f_X_* on two coarse-grained proteome quantities and tested all pairwise combinations of the quantities *e*, *r, r_a_* and 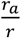 (Supplementary Figure 5). We found that when *f_X_* depends on both the *E* sector concentration *e* and the fraction of active ribosomes 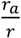, the observed relationship between size and growth rate for the three types of growth rate *r* modulation can be quantitatively captured (Figure 3C and 3E). Notably, the fitted exponent values are very close to 1 and −2/3 for *e* and 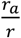 respectively. Because *f_X_* and cell size are inversely proportional, it follows that cell size *V* scales with 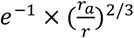. Based on this we propose a novel size law formalized as follows:

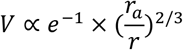

Our model assumes that in the absence of translation inhibition all ribosomes remain active 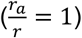. However, it has been observed that for nutrient-limited slow growth the active ribosome fraction is reduced (Dai et al., 2017). Interestingly, incorporating this data in the size law improves its accuracy (Supplementary Figure 6), providing independent support for the proposed relationship between size and active ribosome fraction. Also, the duration of the *C+D* period has been observed to have non-trivial variations (Si et al., 2017). We note that this size law can predict the duration of the *C*+*D* period when incorporating that DNA replication occurs at a fixed condition-independent cell size (Figure 3F).

In summary, our minimalistic model quantitatively captures how both size and coarse-grained cell composition change with the growth rate for all three types of growth modulation with only two parameters (the two exponents of the *f_X_* dependency on *e* and *r_a_*/*r*, those exponents being very close to 1 and −2/3 respectively). Therefore, *E. coli* cell size can be predicted from only two coarse-grained proteome quantities, metabolism sector concentration and the fraction of active ribosomes leading to the definition of a novel growth law.

### Emergence of ‘adder’ size homeostasis and cellular individuality in the presence of noise

So far, we have neglected cell-to-cell variability and focused on predicting average cell composition, size and growth rate at steady-state as a function of growth conditions. However, isogenic cells growing in a constant environment show significant phenotypic variability, notably for global traits such as growth rate or division size (Elowitz, 2002; Levine and Hwa, 2007; Taheri-Araghi et al., 2015). Moreover, non-trivial correlations between single-cell traits are often observed and contain rich information about regulatory mechanisms (Dunlop et al., 2008; Amir, 2014; Kiviet et al., 2014; Taheri-Araghi et al., 2015; Kennard et al., 2016; Kleijn et al., 2018; Thomas et al., 2018; Pandey et al., 2018). Since our model is based on biochemical reactions between coarse-grained molecular components, dynamic cell-to-cell variability naturally emerges when adopting a stochastic interpretation of all reactions and random partitioning of components at division (Figure 4A).

**Figure 4.**
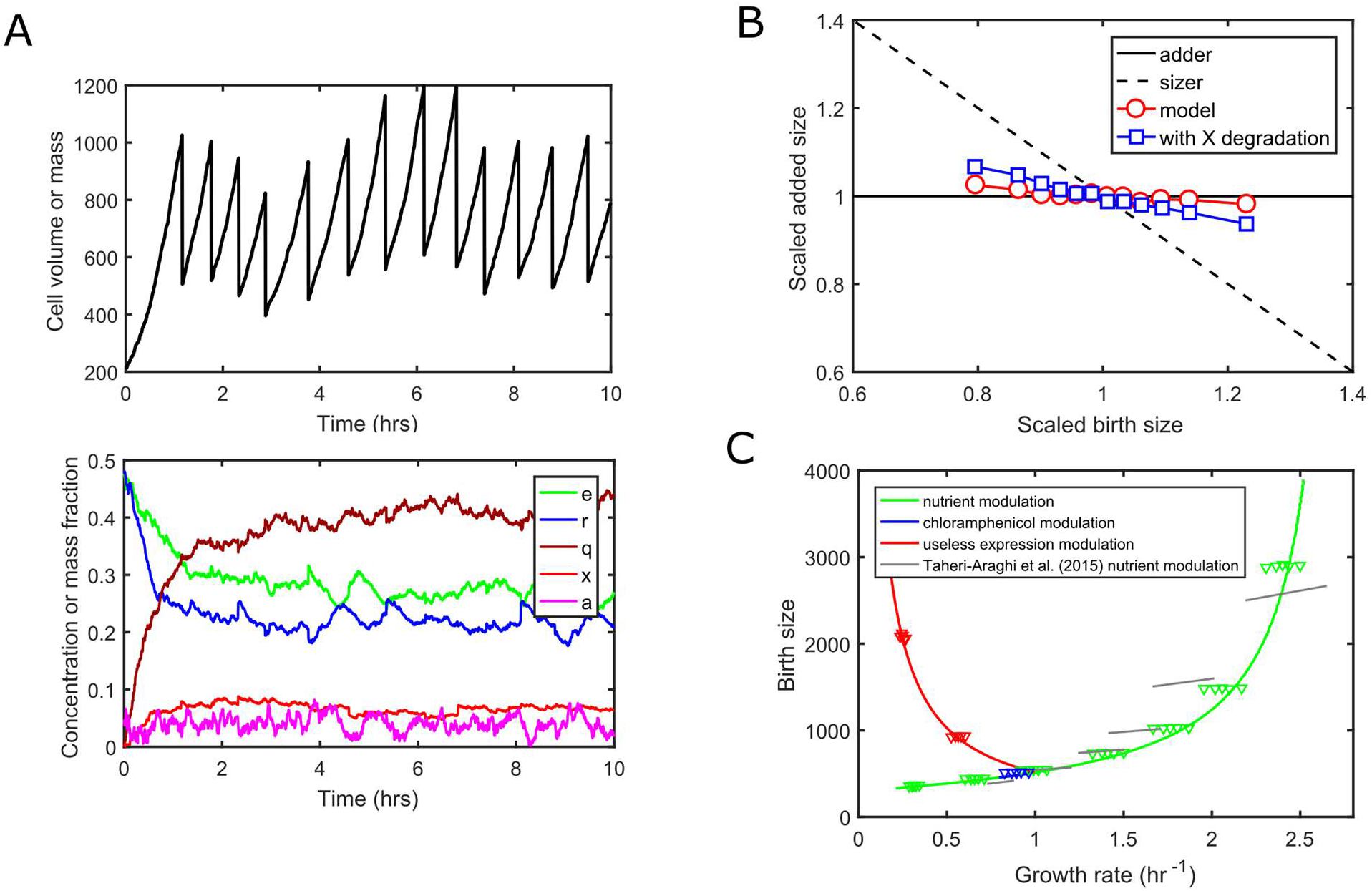
Emergence of ‘adder’ size homeostasis and cellular individuality in the presence of reaction noise. (A) Simulation of model dynamics with molecular noise (top: total cell size, bottom: concentrations of cellular components). At division, cellular components are randomly split between daughter cells, so the tracked daughter has a probability 1/2 of getting each mother cell component. Same parameters as in Figure 2A. The parameters X_div_ and 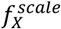 were chosen to obtain realistic cell-to-cell variability in size at birth and growth rate (Methods). (B) Model leads to near ‘adder’ size homeostasis. Average added size during one cell cycle as a function of birth cell size (via binning) is plotted. The very weak deviation towards ‘sizer’ can be explained by a residual correlation between size at birth and X count at birth due to the non-zero share of X in total size. A model variant where X is actively degraded at a constant rate and displaying a stronger deviation towards ‘sizer’ behavior is also shown. Other model variants exploring size homeostasis properties are discussed in Supplementary Figure 7. (C) Emergence of cellular individuality. Stochastic stimulations are performed for ten different growth conditions: seven nutrient qualities (green triangle groups), two useless expression strengths (red triangle groups) and one chloramphenicol (blue triangle groups). For each condition, cell cycles are binned by growth rate and the corresponding birth size is plotted. Continuous lines show the prediction of the deterministic model for the three growth rate modulations. Grey lines indicate experimental trends for different nutrient qualities extracted from mother machine data (Taheri-Araghi et al., 2015).

In the deterministic model, the two parameters *X_div_* (the number of division molecules *X* needed to trigger cell division) and 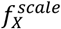 (the constant factor in 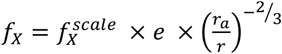) are only setting the size scale via their ratio (Figure 3A). In the presence of reaction noise, those parameters will now impact the extent of cell-to-cell variability in growth rate and in size at division. Even with this simple model of gene expression noise, we could find values of those two parameters that lead to reasonable estimates of both noise in growth rate and size at birth (Methods).

Stochastic simulations of our model lead to a near ‘adder’ size homeostasis, as observed experimentally (Figure 4B, red circles). Interestingly, the deviation towards a ‘sizer’ size homeostasis observed experimentally at very slow growth, (Wallden et al., 2016) could also be explained by additional model assumptions (Supplementary Figure 7). These include the constant degradation of *X* during the cell cycle, which could dominate dilution at slow growth (Figure 4B, blue squares). We then explored the relationship between the size at birth and the growth rate of single cells for several conditions across the three types of growth rate modulation (Figure 4C, triangles). This relationship at the level of single cells is known to deviate from the one connecting average size at birth and average growth rate when conditions are varied (Kennard et al., 2016; Taheri-Araghi et al., 2015). These deviations were qualitatively captured by our stochastic model (Figure 4C, compare grey lines and green triangle groups).

In summary, our minimalistic coarse-grained model generates realistic cell-to-cell variability in growth rate and cell size, as well as realistic correlations between added size and birth size (‘adder’ size homeostasis) and between growth rate and birth size.

## Discussion

In this work, we have proposed a minimalistic whole-cell coarse-grained model of *E. coli* physiology to predict the relationship between growth rate, cell composition, ribosome activity and cell size for three different types of growth rate modulation (Figure 1A and Figure 2). Our model builds on and unifies previous efforts to understand proteome allocation (Molenaar et al., 2009; Scott et al., 2010, 2014; Weiße et al., 2015; Pandey and Jain, 2016) and division control (Fantes et al., 1975; Basan et al., 2015; Taheri-Araghi et al., 2015; Ghusinga et al., 2016). While other theoretical models (Scott et al., 2010, 2014; Weiße et al., 2015; Pandey and Jain, 2016) have also captured proteome allocation data, our model stands out because: 1) it includes ribosome activity and captures more data; 2) it is holistic with respect to cell composition and cell size and does not a priori neglect the mass fraction of protein precursors; and 3) it is not based on optimization of growth rate but rather a positive regulation of ribosome synthesis by precursors. Moreover, despite its simplicity and low number of free parameters (only the exponents in the size law, close to −1 and 2/3 respectively), our model quantitatively reproduces experimental data on proteome allocation Figure (1C-D), ribosome activity (Figure 1E-F) and average cell size (Figure 3C, E) for the three types of growth rate modulation (change of nutrient quality, chloramphenicol-mediated ribosome inactivation and expression of useless proteins). In addition, when reaction noise is included in the model, experimental observations such as the *adder* principle of size homeostasis and non-trivial correlations between single-cell cellular growth rate and cell size are predicted (Figure 4).

A remarkable result of our study is the emergence of a novel size law related to the second growth law. This law states that only one coarse-grained quantity characterizing cell composition is sufficient to predict quantitatively cell size changes in response to nutrient quality and to over-expression of a useless protein (Figure 3C and 3E, Supplementary Figure 4). A second coarse-grained quantity is required for prediction of cell size when translation is inhibited. Those quantities are the mass fraction (or equivalently, concentration) of metabolic proteins (*e* in the model) and the fraction of total ribosomes that are active (*r_a_*/*r*), respectively. Strikingly, best fit is robustly obtained for exponent values close to ~ –1 for the metabolic protein concentration and ~ 2/3 for the fraction of active ribosomes. This results in a novel ‘size law’ stating: 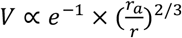. We did not impose constraints on the values of exponents when fitting size data, the fact that fitted exponents are close to such particular values is therefore remarkable and suggests a fundamental underlying physical mechanism. In the absence of translation inhibition and at intermediate to fast growth the fraction of active ribosomes is close to 1 (Dai et al. 2016), therefore the law reduces to the simple form *V* ∝ *e*^−1^. This has been observed before for modulation of growth in the absence of translation inhibition (Taheri-Araghi et al., 2015). Moreover, Basan and colleagues (Basan et al., 2015) noted that if the growth rate dependence of *X* protein follows the one of constitutively expressed proteins for modulation by either nutrient quality or over-expression of useless proteins, then their size data could be explained. Because expression of the metabolic sector has the same growth rate dependence as constitutive expression upon nutrient modulation (Scott et al., 2010), our results agree with those previous observations.

Our ‘size law’ is very different in nature from the second growth law stating that *V* ∝ exp[*α*(*C* + *D*)], where *α* is the growth rate, *C* the duration of DNA replication and *D* the duration between end of DNA replication and cell division. By construction, the second growth law links size at division to the control of DNA replication by size (Donachie, 1968; Wallden et al., 2016; Amir, 2017). It predicts a simple exponential dependency with growth rate for nutrient quality modulation because those three quantities are largely independent of growth rate. However, while average size per origin at replication initiation is invariant across a wide range of growth conditions, both *C* and *D* durations change with growth conditions in complex ways (Si et al., 2017). Therefore, the second growth law is replication-initiation-centric and of limited use to make predictions on cell size at division outside of the canonical type of growth rate modulation (change in nutrient quality). In contrast, the ‘size law’ we propose directly links coarse-grained cell composition to cell size and it explains the observed variation in the *C* and *D* durations at least across three different types of growth rate modulation. This law can generate directly testable predictions, such as cell size and *C*+*D* durations for growth modulations involving both over-expression of useless proteins and translation inhibition by chloramphenicol.

In agreement with our results, recent studies contradict the replication-initiation-centric view of cell size control (Amir, 2017). Based on single-cell correlation statistics, Micali and colleagues (Micali et al., 2018) propose that two concurrent processes, each controlling DNA replication and division, co-determine the effective size at division. Furthermore, Si and colleagues (Si et al., 2019) experimentally demonstrated that two independent ‘adder’ homeostasis mechanisms are in place for DNA replication initiation and cell division. This does not mean that cellular constraints related to DNA replication are not affecting average cell size at division. Yet it indicates that cell division is not enslaved to the process controlling DNA replication initiation. Also, recent experiments reported different amounts of DNA in newborn cells, suggesting the chromosome is not involved in size homeostasis or responsible for the adder behavior (Huls et al., 2018). Here we find that homeostatic properties at the level of cell division appear consistent with the structural model of division control. As shown in Figure 4, the structural model implemented at the single cell level recovers ‘adder’ size control (Ghusinga et al., 2016), deviation towards ‘sizer’ (Wallden et al., 2016) in some parameter regimes (Supplementary Figure 7) and non-trivial deviations from growth law at the single cell level (Kennard et al., 2016; Taheri-Araghi et al., 2015). We note that as suggested by a recent model (Micali et al., 2018), there could be still a role for DNA-replication or segregation in division control, for example when these processes are slowed down (Deforet et al., 2015; Si et al., 2017).

In our model, the quantitative relationship between cell composition and cell size is explained via a dependence of the allocation fraction of division proteins in *e* and 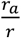 (Figure 3A), but this relationship is also valid in itself as a phenomenological observation (Figure 3C). Thus, the observed growth law is independent of the validity of the structural model and our whole-cell coarse-grained model. What could explain the quantitative relationship between the coarse-grained quantities *e* and 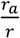 and cell size? Under *r* the structural model of division control, assuming division threshold *X_div_* is invariant across conditions (Figure 3A), it would imply that the *X* allocation fraction *f_X_* scales with *e* and the inverse of 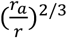. In such case, the dependency on the active ribosome fraction could for example reflect chloramphenicol-mediated alteration of *X* expression, such as operon polarity via premature transcription termination (Zhu et al., 2019). However, under the same assumption that *X_div_* is invariant, Basan and colleagues (Basan et al., 2015) could not identify in proteomics data candidate proteins whose proteome fraction is behaving quantitatively as you would expect for the factor *X,* that is as the inverse of their cell size data. This result suggests that there may be no single protein triggering division when reaching a fixed, condition-independent amount. An alternative possibility consistent with our results is that *X* follows exactly the relative abundance of metabolic proteins *E,* while the division threshold *X_div_* scales with the fraction of active ribosomes to the power 2/3. Other constraints could explain a condition-dependent threshold *X_div_* such as the cell geometry (Harris and Theriot, 2016). A recent study reported that rod-shaped bacteria like *E. coli* maintain an approximately constant aspect ratio across various growth conditions, resulting in the cell surface area scaling as the cell volume to the power 2/3 (Ojkic et al., 2019). It is also possible that the resource allocation that maximizes growth imposes a constraint on the surface-to-volume ratio (i.e., cell width for rod-shaped organisms) between surface proteins and cytoplasmic proteins. In turn, depending on the mechanism “counting” the absolute amount of *X* molecules, the threshold may depend on cell width.

It is also possible that the resource allocation that maximizes growth in a given condition imposes a ratio between surface and cytoplasmic proteins that constrains the surface-to-volume ratio (i.e. the cell width for rod-shaped organisms). In turn, depending on the mechanism ‘counting’ the absolute amount of *X* molecules, the threshold can depend on cell width. In the context of a width-dependent threshold, as suggested by several studies *FtsZ* is an attractive candidate as the sole *X* factor (Palacios et al., 1996; Harris and Theriot, 2016; Zheng et al., 2016; Si et al., 2019). Another mechanism involving geometrical constraints is the recent observation that ribosomes are spatially organized via nucleoid exclusion and that nucleoid size scales with cell size (Gray et al., 2019).

In this work, we have tried to explain the mechanistic origin of size regulation across growth conditions. Whether this is an evolved trait that optimizes fitness is a more difficult question. While size changes due to expression of useless proteins or translation inhibition by drugs may not be an evolved trait, the increased size with nutrient modulation of growth rate seems to be a universal property of microbes, which makes it likely to be an evolved property of unicellular systems (Chien et al., 2012; Turner et al., 2012). In a recent *in silico* study, we showed that increased size at fast growth prevents an increase in gene expression noise for proteins with decreasing concentrations (Bertaux et al., 2018). Metabolic proteins *E* are such proteins, and noise in their expression is likely detrimental to fitness. Interestingly, the inverse relationship between cell size and metabolic protein concentration *e* found in this study implies that the total number of metabolic proteins is kept constant across growth conditions. This raises the possibility that the evolution of size regulation in response to growth conditions has been driven by selective pressures acting on metabolic protein expression noise rather than on size directly. Testing fitness benefit of size regulation is difficult (Monds et al., 2014; Zheng et al., 2016) but should ultimately shed light on the constraints that shaped its evolution.

Modelling cellular systems by considering the cellular context within which they operate is fundamental to a full mechanistic understanding of their function. Modelling has shown that function of simple synthetic networks can be affected by cell cycle and cell growth, when cell physiology and resource allocation is taken into account (Klumpp and Hwa, 2014; Weiße et al., 2015; Borkowski et al., 2016; Bertaux et al., 2018). The current study illustrates that integrating coarse-grained models of cell physiology with simple models of division control could reveal novel size law providing a link between cell size and cellular resource allocation. Overall, our study highlights the importance of systematic integration of whole-cell physiological models in the study of natural and synthetic cellular systems.

## Acknowledgements

We would like to thank Suckjoon Jun, Istvan Kleijn, Guillaume Le Treut, Peter Sarkies, Fangwei Si, Peter Swain and Philipp Thomas for feedback on our manuscript and members of the Marguerat and Shahrezaei groups for discussions. F.B. received financial support from Leverhulme Research Project Grant (RPG-2014-408) awarded to S.M. and V.S. V.S. is supported by the EPSRC Centre for Mathematics of Precision Healthcare (EP/N014529/1). S.M. is supported by the UK Medical Research Council.

## Methods

### Model reactions

Our model considers the following reactions between coarse-grained cell components:

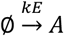 (nutrient import and transformation into protein precursors A)
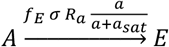 (synthesis of *E* proteins)
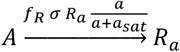 (synthesis of active *R* proteins)
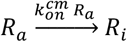 (inactivation of active *R* proteins via chloramphenicol binding)
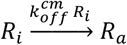 (re-activation of chloramphenicol-inactivated *R* proteins via chloramphenicol unbinding)
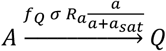 (synthesis of *Q* proteins)
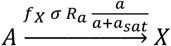 (synthesis of *X* proteins)
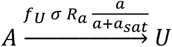 (synthesis of *U* proteins)

Where the cell volume is *V* = *E* + *R* + *Q* + *X* + *U* + *A* (with *R* = *R_a_* + *R_i_*), the protein synthesis allocation parameters (*f_E_*, *f_R_*,…) respect the constraint *f_E_* + *f_R_* + *f_Q_* + *f_X_* + *f_U_* = 1, and 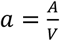(we will also note *e, r, q, x, u*,… the concentrations *E/V, R/V*, etc…). The parameter *k* is the medium nutrient quality, 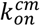 is the chloramphenicol-imposed inactivation rate of ribosomes and *f_U_* is the imposed allocation fraction of useless proteins. Those three parameters represent the three types of growth rate modulations that we consider. The parameter *σ* is the maximal rate of protein synthesis by one ribosome and *a_sat_* is the enzymatic saturation constant of ribosomes. The parameter 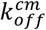 is the dissociation rate of chloramphenicol-ribosome complexes. Note that all protein synthesis reactions conserve mass, so that volume growth equals nutrient import (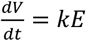 as seen below).

### Differential equations

A deterministic interpretation of these reactions leads to the following set of differential equations for absolute amounts of coarse-grained cell components:

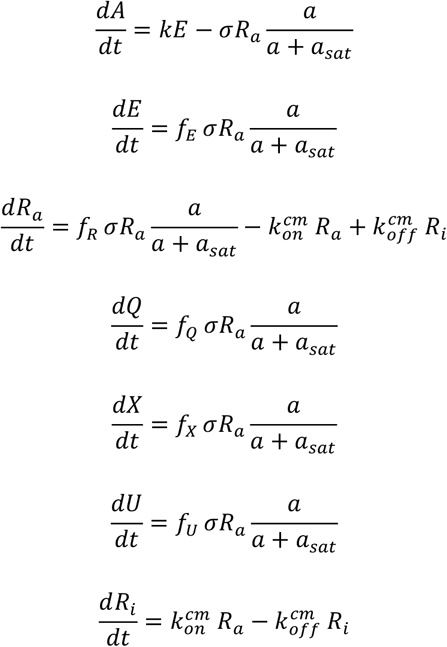

The above differential equations can be also written in terms of concentrations, noticing that 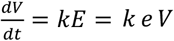:

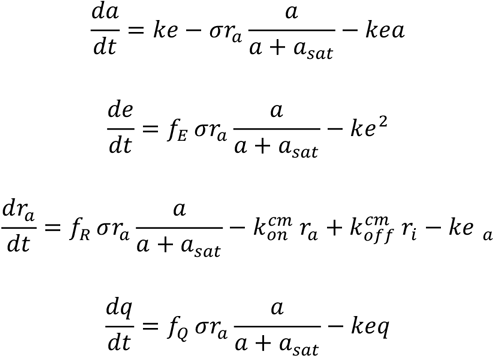

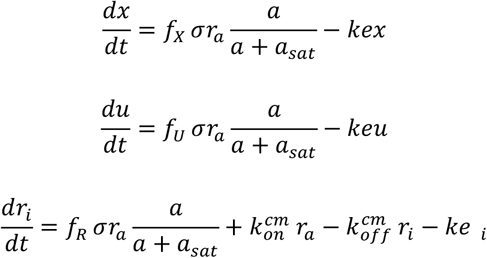

In the above equations, the last terms represent dilution by cell growth (*ke*). Because concentrations are conserved at division, their dynamics and steady-state values are independent of the division threshold *X_div_*.

### Steady-state growth

In this model, steady-state growth (or balanced growth, i.e. steady concentrations of coarse-grained cell components) is necessarily exponential at rate *α* = *k e*, because 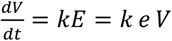. By setting the time derivatives to zero, we obtain the following equations constraining the steady-state cell composition (*e, r_a_, r_i_, q, x, u, a*):

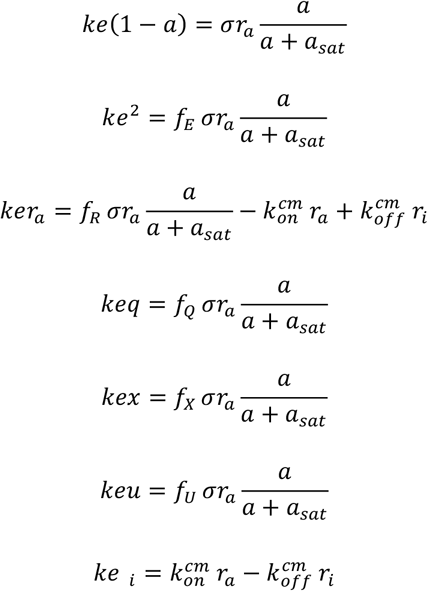

The first equation gives another expression for the growth rate: 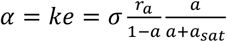, where the factor 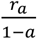 is in fact the proteome fraction of active ribosomes *φ_Ra_*. It also reflects the balance between the rate of nutrient import and transformation into precursors and the rate of total protein synthesis in the context of dilution imposed by mass density homeostasis.

Combining the first equation with every other enables expressing protein concentrations as a function of (1 – *a*):

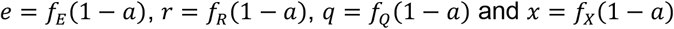

This is intuitive: a proteome sector concentration is equal to its allocation fraction times the total protein concentration 1 – *a*. We also see that steady-state proteome fractions *φ* are equal to allocation fractions *f*.

Noting that 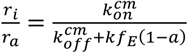 and that 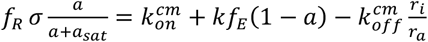 leads to an equation for 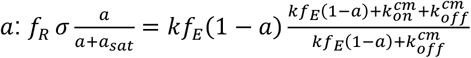

Solving this equation for *a* allows to compute the steady-state solution for fixed allocation fractions *f_R_*, *f_Q_*, *f_X_* and *f_U_*.

### Dynamic regulation of allocation fractions

Dai and colleagues (Dai et al. 2016) found that the translation elongation rate displays a Michaelis-Menten like dependence with the RNA to protein ratio (which is proportional to the total ribosome proteome fraction) across nutrient conditions and chloramphenicol-mediated translation inhibition. A simple way to reproduce these findings with our model is to assume that the allocation fraction *f_R_* is regulated by the concentration of precursors via direct proportionality: *f_R_* = *δ* · *a*. Indeed, we would then have 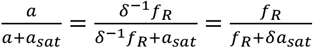 Note that such regulation is a form of supply-driven activation (Scott et al. 2015).

Therefore, our model for how allocations fractions are dynamically set is as follows:

1. *f_Q_*, *f_U_*, *f_X_* are constants (*f_Q_* is also invariant across conditions)
2. *f_R_* = *δ* · *a* (or 1 – /– *f_Q_* – *f_U_* – *f_X_*)
3. f_E_ = 1 – *f_R_* – *f_Q_* – *f_U_* – *f_X_*

The differential equations presented above with such dynamically changing allocations fractions are used to simulate Figure 2A, together with the halving of all amounts at division, which is triggered when *x* = *X_div_*.

### Steady-state proteome fractions for the regulation model

By substituting *f_R_* = *δ* · *a* in the steady-state growth equations above, we obtain the following equation if we further assume that *a* ≪ 1 (i.e. *δ* is large enough):

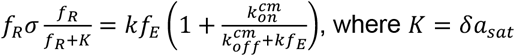

Because *f_E_* = 1 – *f_R_* – *f_Q_* – *f_U_* – *f_X_*, this can be solved numerically by scalar optimization on *f_R_*.

This steady-state approximation was used for the results presented in Figure 1 and 3. However, when simulating the dynamic model (Figure 2 and 4), a value for *δ* has to be chosen, and the allocation fraction *f_X_* has to be non-zero. We chose *δ* = 5 and we subtracted *f_X_* from *f_Q_* to ensure *f_Q_* + *f_X_* = 0.5. We show in Supplementary Figure 1 the validity of the *a* ≪ 1 approximation.

### Optimality of proteome allocation fractions

Above we considered the case where the allocation fractions are dictated by a supply-driven activation of ribosome synthesis. For given growth conditions, one could also search for the proteome allocation fractions that maximize growth rate (under the constraints of a fixed, condition-independent allocation fraction to housekeeping proteins *f_Q_*, an imposed allocation fraction to useless proteins *f_U_*, and under the assumption that *f_X_* ≪ 1).

This is achieved by enumerating all possible allocations by varying *f_R_* in between 0 and 1 – *f_Q_* – *f_U_*, and computing the growth rate of the corresponding steady-state (we use the bounded scalar optimization *MATLAB* function *fminbnd*).

Such optimality assumptions are sometimes made in previous work, such as in Molenaar et al. (2009).

### Model parameterization

Cell composition data is taken from (Scott et al. 2010) and (Dai et al. 2016). It consists of steady-state ribosome proteome fractions for a combination of nutrient conditions and chloramphenicol concentrations (measured via the RNA/protein ratio). The dataset from Dai et al. (2016) also provides translation elongation rate and estimated fraction of ribosomes that are active. Following Scott and colleagues (Scott et al., 2010), we used a conversion factor of 0.76 to convert RNA/protein ratios into proteome fractions of extended ribosomes. For consistency, we also used the notion of extended ribosomes (1.67 times larger than a single ribosome, Scott et al., 2010) to convert the maximal elongation rate of 22 aa/s measured by Dai and colleagues (Dai et al., 2017) into a maximal rate of protein synthesis (*σ* in our model). Finally, based on maximal ribosome proteome fractions observed by Scott and colleagues, we set the value of *f_Q_* to 0.5. Altogether, we obtained the following parameterization from those studies:

- 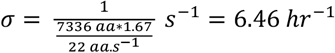
- *K* = 0.11 * 0.76 = 0.0836
- *f_Q_* = 0.5

### Finding regulation of *f_X_* expression explaining cell size data across types of growth rate modulation

The structural model of cell division links the steady-state cell composition and cell division size via the concentration *x* of *X* proteins: 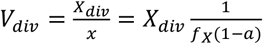, which approximates to 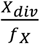 under the low precursor concentration assumption.

To search for regulation of *f_X_* explaining cell size data across the three types of growth rate modulations, we considered the following form of 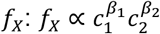 where *c_1;2_ are quantities depending on the coarsegrained cell composition and *β**_1,2_ are exponents.

We considered the following coarse-grained quantities: coarse-grained concentrations *e*, *r*, *r_a_* and the fraction of active ribosomes 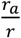 (not to confuse with the mass fraction of active ribosomes *r_a_* or the proteome fraction of active ribosomes 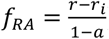).

Fitting was performed using multilinear regression on the logarithm of cell size data using *MATLAB* function *regress*. 95% confidence intervals on exponents were provided by the *regress* function. Cell size data between different datasets was normalized as detailed in Supplementary Figure 3.

To estimate the *C*+*D* duration directly from our size estimate, we use the value of ‘unit cell’ or DNA replication initiation volume from (Si et al. 2017) (*S*_0_ = 0.28 *μm*^3^, converted to the normalized size scale described in Supplementary Figure 3).

### Stochastic model

The stochastic version of the model was simulated using the *Gillespie* algorithm and in a mother machine setting. When *X* reaches the division threshold *X_div_*, each coarse-grained molecule is kept in the daughter cell that we keep simulating with probability ½. Note that we are modeling protein synthesis with a single reaction for each sector, hence abstracting away transcription and mRNA degradation.

### Code availability

All the code for solving model steady-state, simulating model dynamics, parameter fitting, etc. as well as the scripts generating all figures is available on GitHub. It is written in *MATLAB* except the stochastic simulation code written in *C*++.

## Notes

#### Summary of Updates

The model now captures recent ribosome activity data (Dai et al., 2017) in addition to proteome allocation data, size and growth rate data.

