## Supplementary Figures for "A bacterial size law revealed by a coarse-grained model of cell physiology"

1. Department of Mathematics, Imperial College London, London SW7 2AZ, UK
2. MRC London Institute of Medical Sciences (LMS), London W12 0NN, UK
3. Institute of Clinical Sciences (ICS), Faculty of Medicine, Imperial College London, London, W12 0NN, UK
4. Institut Pasteur and CNRS, C3BI - USR 3756, 25–28 Rue du Docteur Roux, 75015, Paris, France
5. To whom correspondence should be addressed

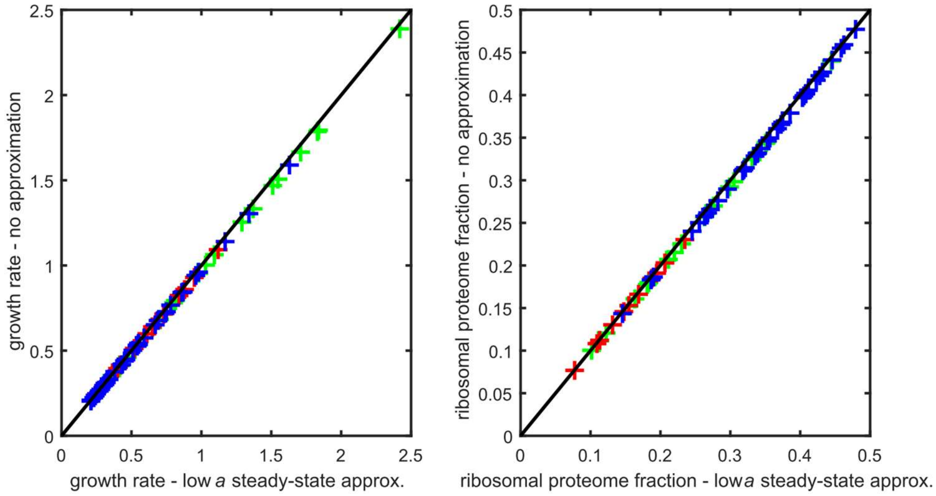


***Supplementary Figure 1. Validity of the low precursor concentration approximation.*** *Comparison of steady-state predictions between the low precursor concentration approximation model and its exact counterpart. Left: growth rate, Right: R proteome fraction* $\varphi_R$*. Each point corresponds to a growth condition (green: nutrient modulation, red: useless expression modulation, blue: chloramphenicol-mediated ribosome inactivation). For both models,* $f_Q = 0.5,\ \sigma = 6.46\,hr^{-1},\ K = 0.0836$*. For the exact model,* $\delta = 5$ *and thus* $a_{sat} = \frac{K}{\delta} = 0.0167$*. For the exact model steady-state were computed by simulating the dynamic model for long enough.*

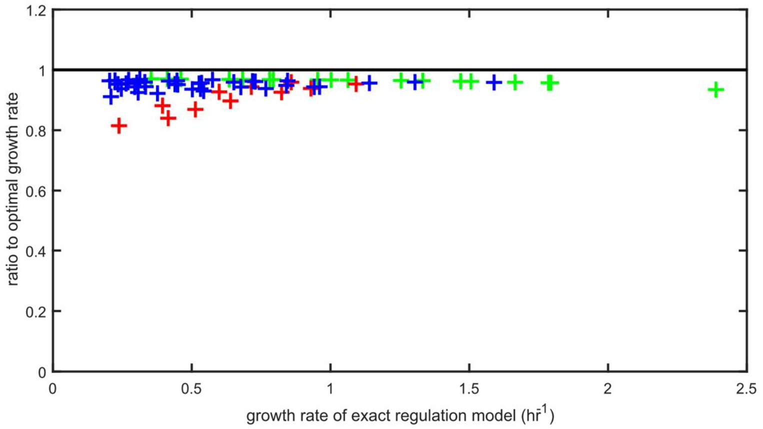


***Supplementary Figure 2. The dynamic regulation model achieves near-optimal proteome allocation across growth conditions.*** *The optimality ratio of the dynamic regulation strategy ($\delta = 5$) is shown as a function of growth rate for a range of growth conditions (green: nutrient modulation, red: useless expression modulation, blue: chloramphenicol-mediated ribosome inactivation). For both models,* $f_Q = 0.5$, $\sigma = 6.46\ hr^{-1}$, $a_{sat} = 0.0167$.

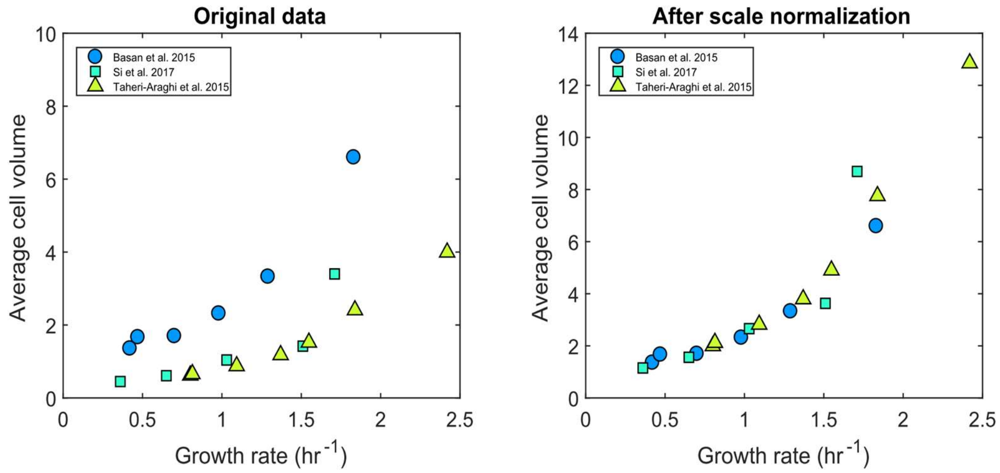


***Supplementary Figure 3. Scale normalization of cell size data.*** *Size - growth rate data for from three different sources (Basan et al., 2015; Si et al., 2017 and Taheri-Araghi et al., 2015) displayed systematic differences for nutrient modulation (left). We searched for scalar correction factors to normalize this data. We found that multiplying Si et al. data by 2.56 and Taheri-Araghi et al. data by 3.22 was sufficient to obtain a consistent size - growth rate relationship (right). Those factors were then applied on all data from those studies (including other types of growth rate modulation) for further analysis.*

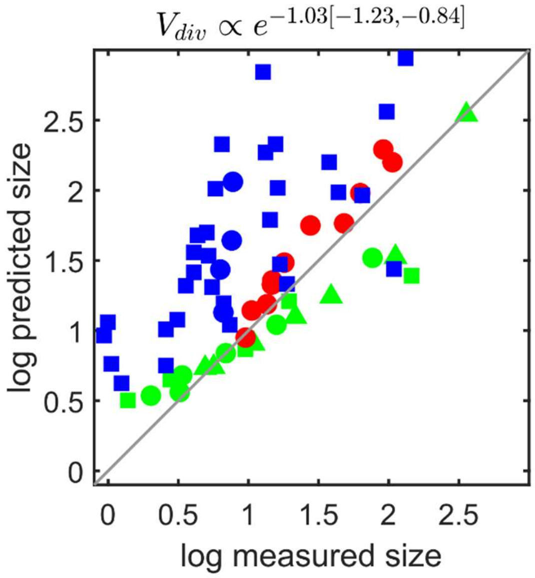


***Supplementary Figure 4. The metabolic sector concentration predicts cell size for nutrient and useless expression modulations but not for chloramphenicol-mediated ribosome inactivation.*** *The size law of Figure 3D is tested against all three growth rate modulations (green: nutrient modulation, red: useless expression modulation, blue: chloramphenicol-mediated ribosome inactivation). Data sources are Basan et al., 2015 (circles), Si et al., 2017 (squares) and Taheri-Araghi et al., 2015 (triangles). For chloramphenicol-mediated ribosome inactivation, cell size is systematically over-estimated by the metabolic sector concentration size law.*

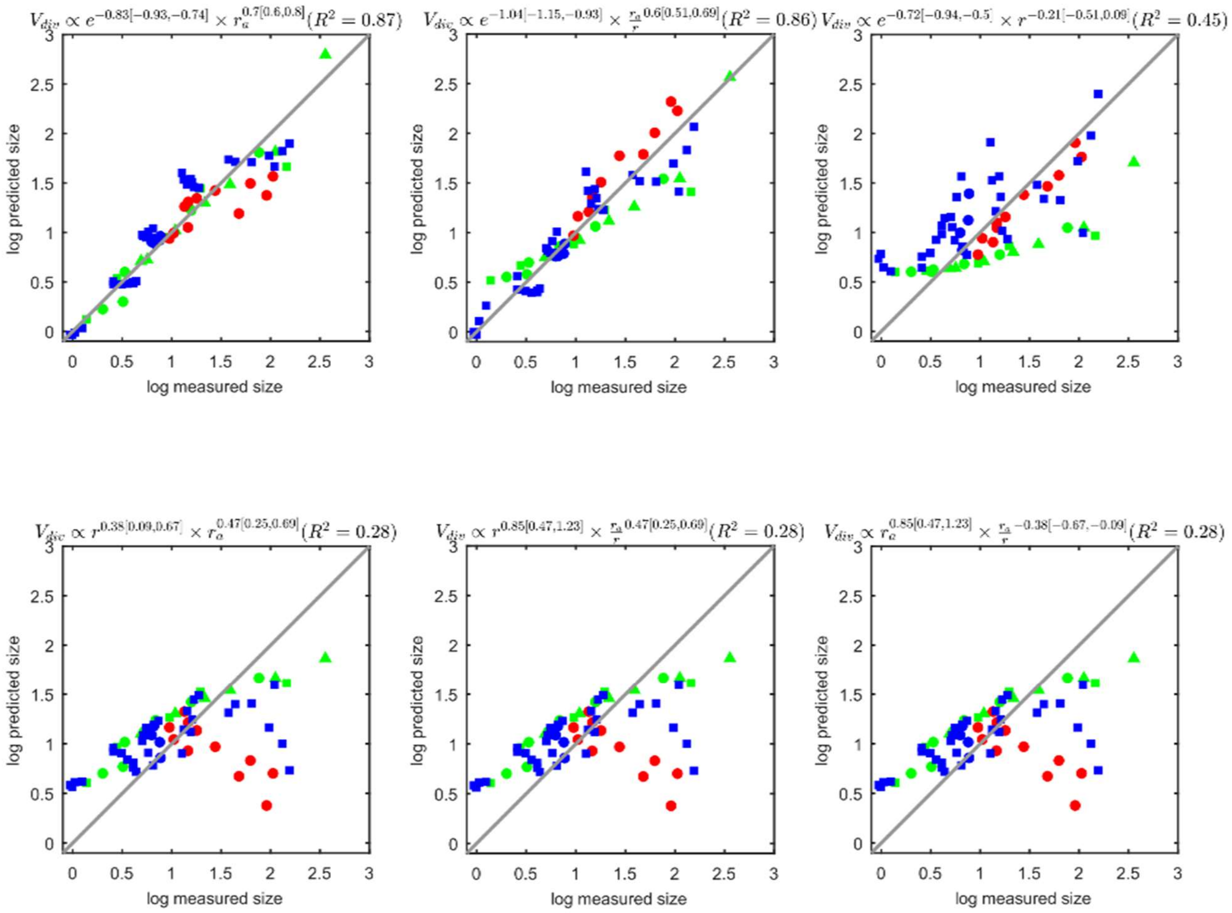


***Supplementary Figure 5. Predictability of cell size from two coarse-grained proteome quantities.*** *The model is used to compute the coarse-grained quantities $e$, $r$, $r_a$ and $\frac{r_a}{r}$ for each growth condition (green: nutrient modulation, red: useless expression modulation, blue: chloramphenicol-mediated ribosome inactivation). Pairwise combinations of those quantities are regressed against cell size in log space. The corresponding best 'size laws' are shown above each plots, together with 95% confidence intervals for the exponents and the $R^2$ of the regression. Data sources are Basan et al., 2015 (circles), Si et al., 2017 (squares) and Taheri-Araghi et al., 2015 (triangles). Note that the three bottom combinations are in fact equivalent.*

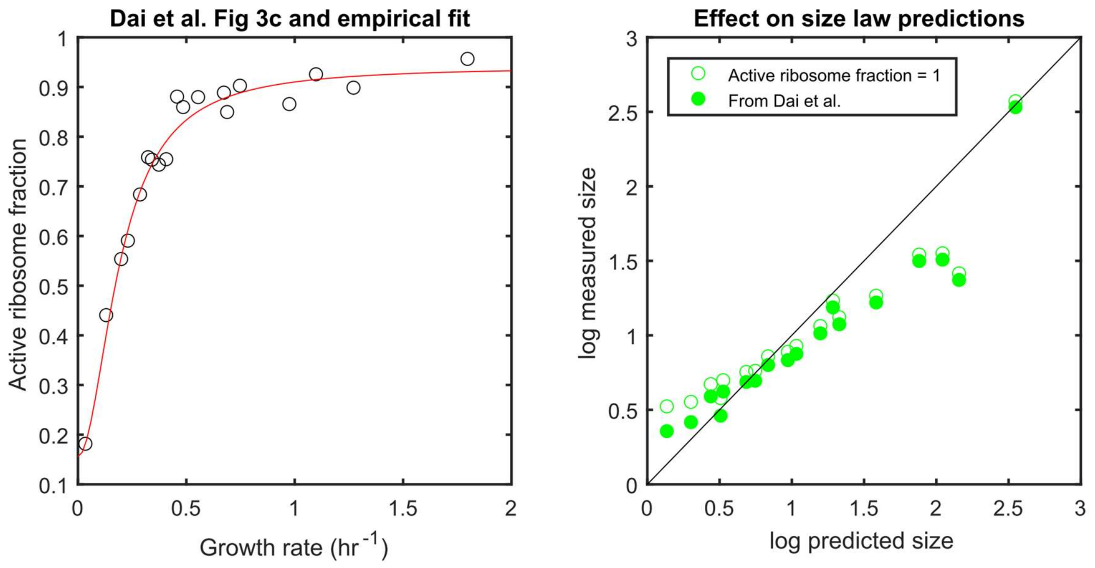


***Supplementary Figure 6. Accounting for inactive ribosomes in poor nutrient conditions improves predictions of the size law.*** *Our model assumes that in absence of chloramphenicol, all ribosome are active, yet at very slow growth high fractions of inactive ribosomes have been found (**left**). The red line is an hill function fit to extrapolate the data to all growth rates. When such growth rate dependent active ribosome fractions are accounted for in the size law of Figure 3, the prediction error at slow growth (small sizes) is decreased **(right)**.*

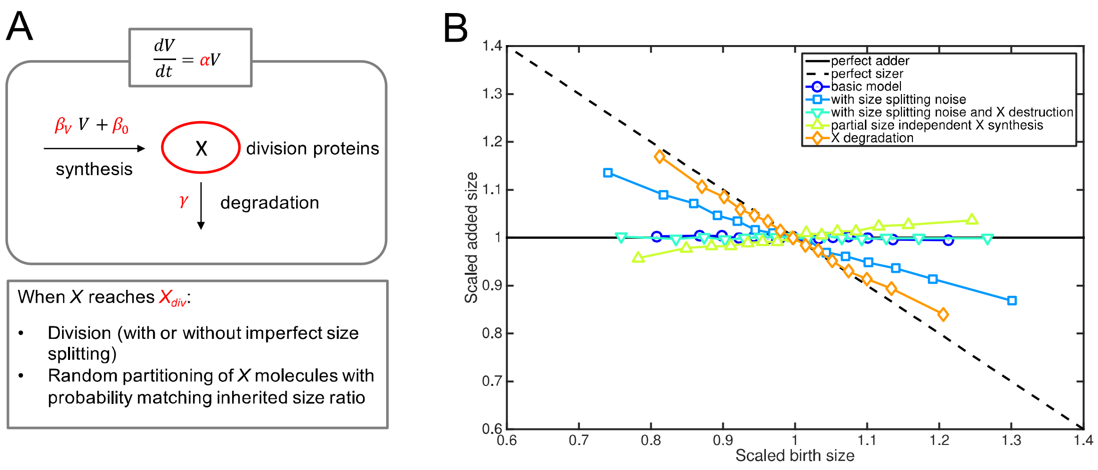


***Supplementary Figure 7. Many simple assumptions can generate deviations from adder size homeostasis within the structural model of cell division.*** *(A) Description of the model structure. In all model variants, cell size is growing exponentially at a constant rate. All model variants lead to stable distributions of size at division. (B) Size homeostasis properties for different model variants. The basic model leads to adder behavior ($X_{div} = 90,\ \alpha = 1\ hr^{-1},\ \beta_V = 30\ hr^{-1}\ per\ size\ unit, \beta_0 = 0\ hr^{-1}$, no size*

*splitting error). When the size splitting noise is high (standard deviation of 0.1 around equal splitting of 0.5), deviation towards sizer is observed. Destroying X at division restores adder behavior despite strong size splitting noise ($\beta_V = 60\ hr^{-1}\ per\ size\ unit$ to obtain the same average division size). If X synthesis is partially size-independent, a small deviation towards an inverse sizer can be observed ($\beta_V = 25\ hr^{-1}\ per\ size\ unit$ and $\beta_0 = 10\ hr^{-1}$, chosen again to obtain the same average division size). Constant first-order degradation of X also generates a deviation towards sizer size homeostasis (here $\gamma = 2\ hr^{-1}$, and $\beta_V = 120\ hr^{-1}\ per\ size\ unit$ to have a comparable average division size).*
